# A genome and tissue-specific transcriptomes of the large-polyp coral, *Fimbriaphyllia* (*Euphyllia*) *ancora*: Recipe for a coral polyp

**DOI:** 10.1101/2023.12.17.572093

**Authors:** Shinya Shikina, Yuki Yoshioka, Yi-Ling Chiu, Taiga Uchida, Emma Chen, Yin-Chu Cheng, Tzu-Chieh Lin, Yu-Ling Chu, Miyuki Kanda, Mayumi Kawamitsu, Manabu Fujie, Takeshi Takeuchi, Yuna Zayasu, Noriyuki Satoh, Chuya Shinzato

## Abstract

Coral “polyps” are composed of several tissues; however, their characteristics are largely unexplored. Here we report biological characteristics of the four tissues that comprise polyps: tentacle (*Te*), mesenterial filament (*Me*), body wall (*Bo)*, and mouth with pharynx (*MP*), using comparative genomic, morpho-histological, and transcriptomic analyses of the large-polyp coral, *Fimbriaphyllia ancora*. A draft *F. ancora* genome assembly of 434 Mbp was created. Morpho-histological and transcriptomic characterization of the four tissues showed that they have distinct differences in structure, primary cellular composition, and transcriptional profiles. Tissue-specific, highly expressed genes (HEGs) of *Te* are related to biological defense, predation, and coral-algal symbiosis. *Me* expresses multiple digestive enzymes, whereas *Bo* expresses innate immunity and biomineralization-related molecules. Many receptors for neuropeptides and neurotransmitters are expressed in *MP*. The established dataset and new insights into tissue functions will facilitate a deeper understanding of coral biology.

## Introduction

Scleractinian corals are the primary builders of coral reef ecosystems, which nurture a wide variety of marine organisms (Odum and Odum, 1955; Fisher et al., 2015). Coral reefs support tropical and sub-tropical fisheries and tourism (De Groot et al., 2012). Despite their ecological and economic importance, biological properties of corals are still largely unknown. A comprehensive understanding of coral biology will not only allow more accurate predictions of anthropogenic impacts on corals (Shinzato et al., 2011), but may help to establish methods for preservation and propagation of extant corals.

Corals belong to the phylum Cnidaria, a group of animals possessing cnidocytes (Ruppert et al., 2003). Generally, corals and other cnidarians, e.g., sea anemone, hydra, and jellyfish, are diploblastic and regarded as animals with simple body structures. Tissues, organs, and organ systems characteristic of vertebrates are not present in corals (Technau and Steele, 2011; Shikina and Chang, 2018). Yet, individual coral “polyps” are composed largely of several functionally defined tissues: tentacle, mouth with pharynx, mesenterial filament, and body wall (Ruppert et al., 2003). Tentacles, located atop each polyp, are responsible for predation, attack, and defense. The mouth with the associated pharynx is located in the upper part of the polyp not only to engulf prey, but also to release excreta, gametes, and larvae. Mesenterial filaments are located in the body cavity and constitute the primary tissue responsible for digestion and defense. The body wall separates the interior of a coral polyp from the environment. Although differences and functions can be inferred from tissue locations and behavioral observations (Nugues et al., 2004; Shikina et al., 2023a), molecular and cellular characteristics and detailed functions of those tissues remain largely unexplored. Fine characterization of each tissue comprising coral polyps will lead to a better understanding of biological properties of corals.

Tissue-specific analysis is a powerful approach to characterize structure, cellular composition, and function. Detailed cellular and molecular analyses of isolated target tissues, rather than whole individuals, allow us to explore differences in organization and function among tissues, as well as intrinsic mechanisms underlying specific functions (Su et al., 2004; Fagerberg et al., 2014). However, tissue-specific analyses have rarely been conducted in corals. The reason for this is that the coral species often used for cellular and/or molecular studies, e.g., *Acropora millepora* (Attenborough et al., 2019), *Pocillopora damicornis* (Domart-Coulon et al., 2001), *Stylophora pistillata* (Muscatine et al., 1997), have very small polyps (about 1 mm in diameter), making tissue isolation difficult. *Fimbriaphyllia ancora* (family Euphyllidae), formerly called *Euphyllia ancora* (**Fig. 1 A**; Luzon et al., 2017), is distributed widely in reefs of the Indo-Pacific Ocean (Luzon et al., 2017; Hoeksema and Cairns, 2023) and is one of the most popular coral species in the aquarium industry (Bruckner, 2001). Morphologically, it has tentacles with swollen, anchor-like tips and a flabello-meandroid skeleton **(Fig. 1 B and C)** (Luzon et al., 2017). The most notable morphological characteristic is its polyp size (3-5 cm in diameter), and techniques for isolating tissues from these polyps have been established (Shikina et al., 2013).

**Figure 1.**
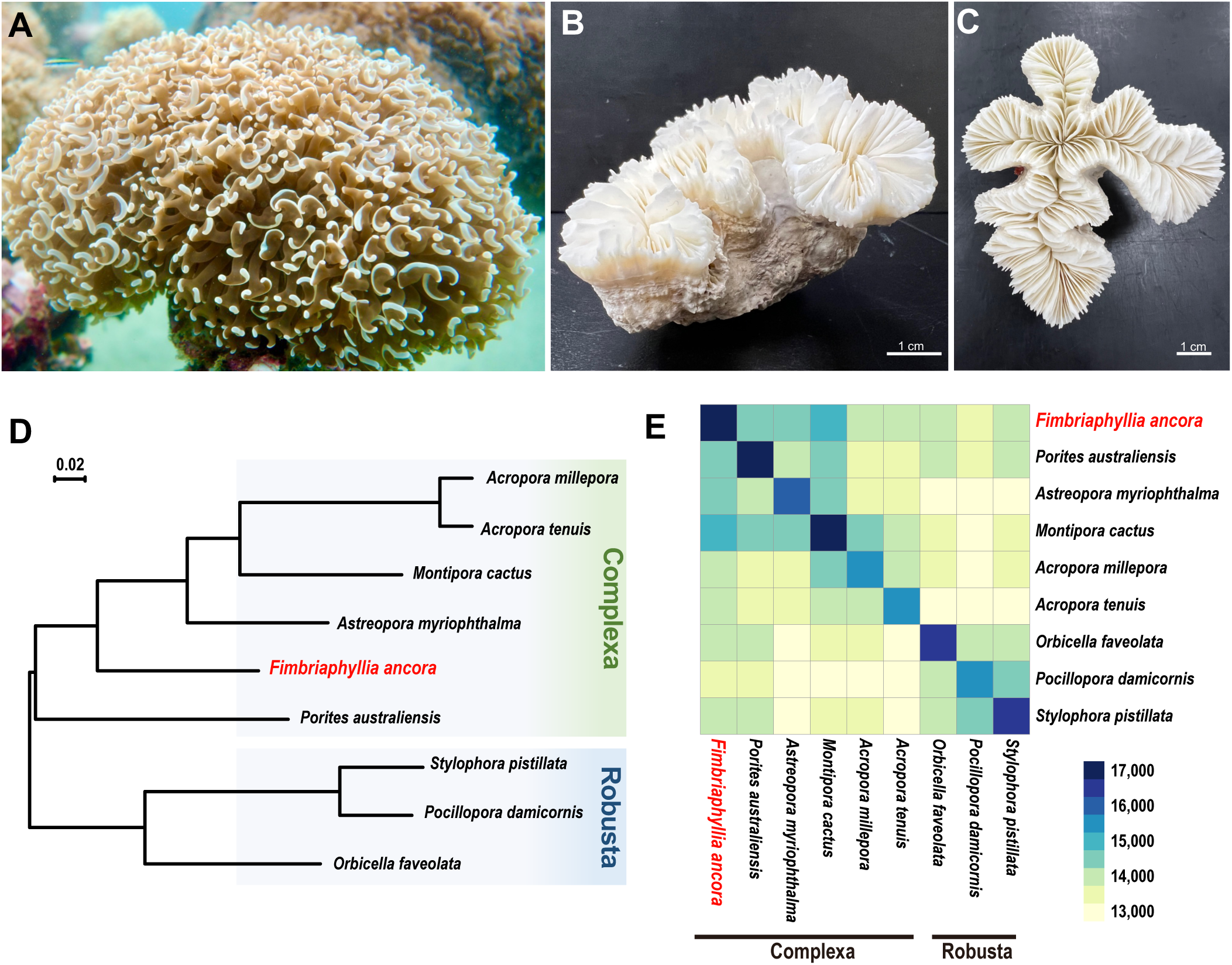
*Fimbriaphyllia ancora* and its phylogenetic relationships with other scleractinian corals. **A**. External appearance of an *F. ancora* colony. **B** and **C**. Top and side views of an *F. ancora* skeleton. The skeleton was photographed after removal of polyp tissue. Hammer or anchor-like tentacles and the flabello-meandroid skeleton typify *F. ancora.* **D**. Molecular phylogeny of *F. ancora* based on 4,208 single-copy Orthogroups (OGs) identified from published scleractinian genomes. All nodes are supported with 100% bootstrap values. **E**. A heatmap showing numbers of shared OGs among scleractinian genomes.

Using *F. ancora*, we sought to determine cellular and molecular characteristics of the four representative tissues (tentacle, mesenterial filament, body wall, and mouth with pharynx) constituting coral polyps. Ultimately, such fundamental data will enhance our understanding of biological characteristics of corals. To this end, we generated a draft genome of *F. ancora* and performed morpho-histological and transcriptomic analyses of all four polyp tissues.

## Results

### The Fimbriaphyllia ancora draft genome

A total 14.6 Gbp of PacBio HiFi reads (QV>20, 1.8M reads, average length: 8,145bp) were obtained. Then, we assembled the reads into an *F. ancora* draft genome assembly of 434 Mbp, comprising 205 scaffold sequences with an N50 size of 5.18 Mbp (Table 1). No scaffold sequences showed significant similarity with reported Symbiodiniaceae genomes (BLASTN, e-value <e^-10^) (**Table S1**), indicating no contamination with dinoflagellate sequences. A total of 27,537 protein-coding genes were predicted, and the gene number was comparable to those of other coral genomes (**Table 1**). Benchmarking Universal Single-Copy Orthologs (BUSCO) analyses (Simão et al., 2015; Waterhouse et al., 2018), which assess whether universal single-copy orthologous genes observed in more than 90% of metazoan species (from the OrthoDB database of orthologs) (www.orthodb.org; version 9) are recovered in a genome/transcriptome assembly, yielded completeness scores for the *F. ancora* genome assembly and gene models of about 95.6% and 96.1% (Complete BUSCO %), respectively. This indicated that the genome assembly and gene models are of comparable quality to those of previously reported coral genomes.

**Table 1.**
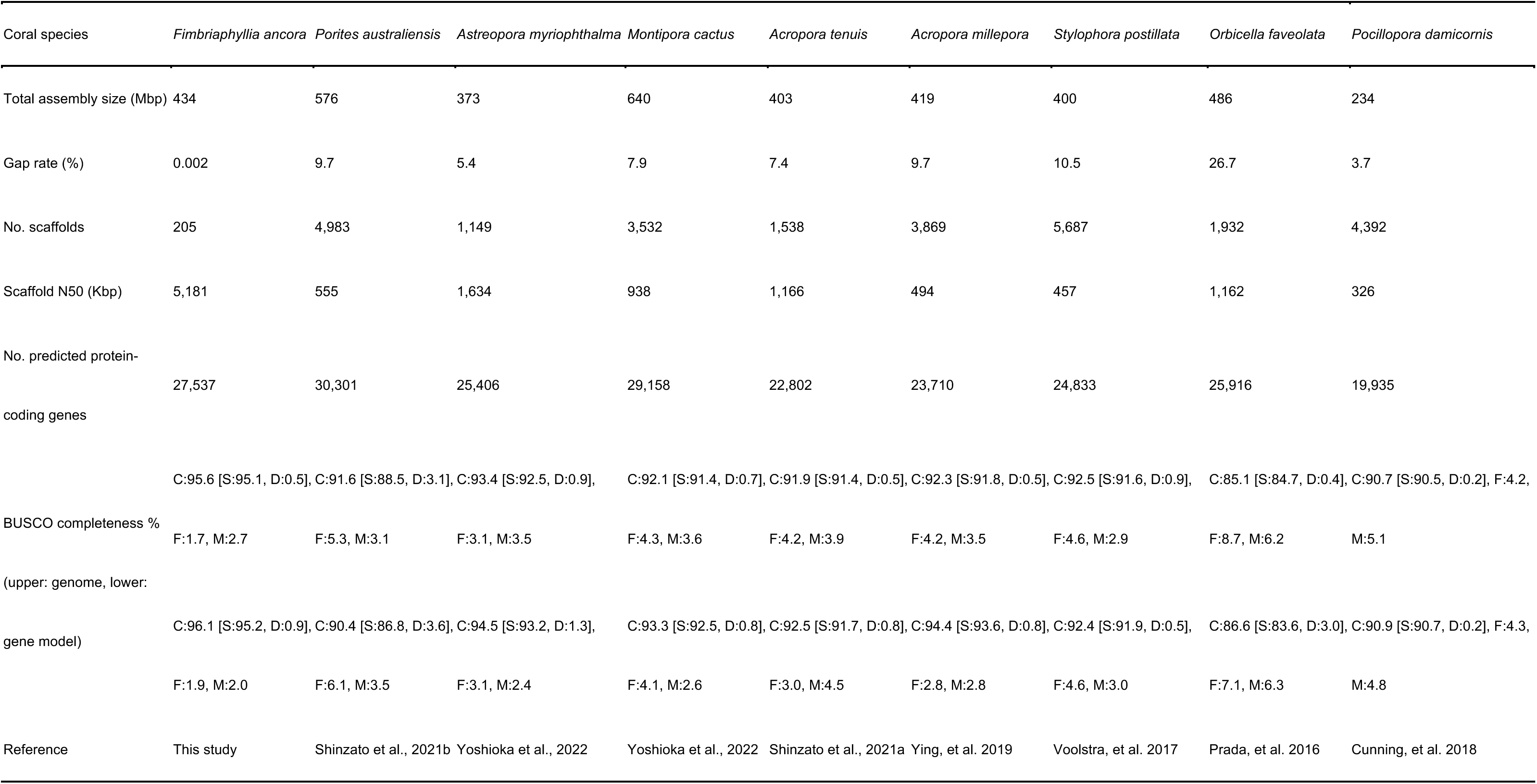
Genome assembly and gene prediction statistics for *Fimbriaphyllia ancora* and comparisons with publicly available scleractinian coral genomes.

Next, orthologous relationships with other scleractinian corals were investigated using genome sequences of 4 acroporid species (*Acropora millepora*, *Acropora tenuis*, *Montipora cactus,* and *Astreopora myriophthalma*), *Porites australiensis*, *Stylophora pistillata*, *Pocillopora damicornis*, and *Orbicella faveolata*. We identified 23,701 orthologous gene families (OGs) from scleractinian genomes and obtained 17,128 OGs for *F. ancora*. Phylogenomic analysis of these anthozoan genomes using concatenated amino acid sequences of 4,208 single-copy orthologous group genes (1,817,638 AAs) yielded robust phylogenetic relationships, with all clades supported by 100% bootstrap values **(Fig. 1 D)**, clearly indicating that *F. ancora* belongs to the Complexa coral clade, as reported by previous molecular phylogenic analyses (Fukami et al., 2008; Kitahara et al., 2010). Among the OGs in *F. ancora*, 10,016 are shared by all scleractinians, 16,511 OGs are shared with the Complexa group, and 371 gene families were restricted to *F. ancora* **(Fig. 1 E)**.

### Morphological and histological characteristics of the four tissues comprising F. ancora polyps

Four major tissues, tentacle, mesenterial filament, body wall, and mouth with pharynx **(Fig. 2 A-C)** were isolated, and their morpho-histological characteristics were examined **(Table 2)**.

**Figure 2.**
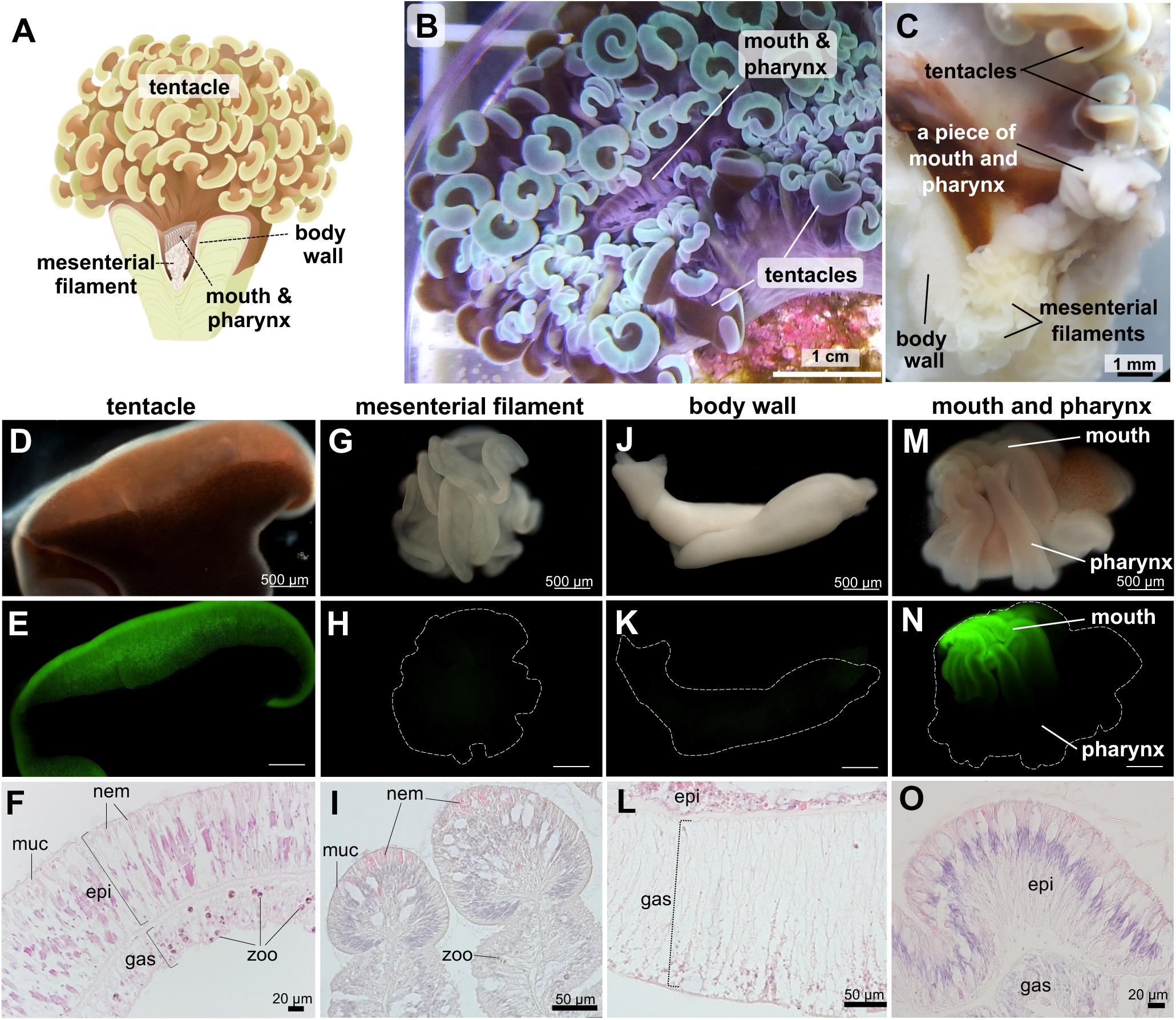
**A.** Schematic diagram of a *Fimbriaphyllia ancora* polyp showing the 4 tissues analyzed in this study. **B**. Top view of an *F. ancora* colony in an aquarium. The mouth is at the center of the polyp, and tentacles surround the mouth. **C**. Representative picture of a dissected *F. ancora* polyp with tentacles, part of the mouth with the pharynx, mesenterial filaments, and part of the body wall. **D-O**. Appearances of isolated polyp tissues and their histological micrographs. **D-F**. A tentacle. **G-I**. A mesenterial filament. **J-L**. A piece of the body wall. **M-O**. A piece of the mouth with the pharynx. **D, G, J, and M**. Bright views of isolated polyp tissues. **E, H, K, and N**. U-MWIB 2 (GFP) filter view of the same field as in **D**, **G**, **J**, and **M**. Dashed lines in **H**, **K**, and **N** show outlines of each tissue. **F, I, L, and O**. Histological sections of isolated tentacle, mesenterial filament, body wall, and mouth and pharynx, respectively. epi, epidermis; gas, gastrodermis; zoo, zooxanthellae; nem, nematocyte; muc, mucocytes.

**Table 2.**
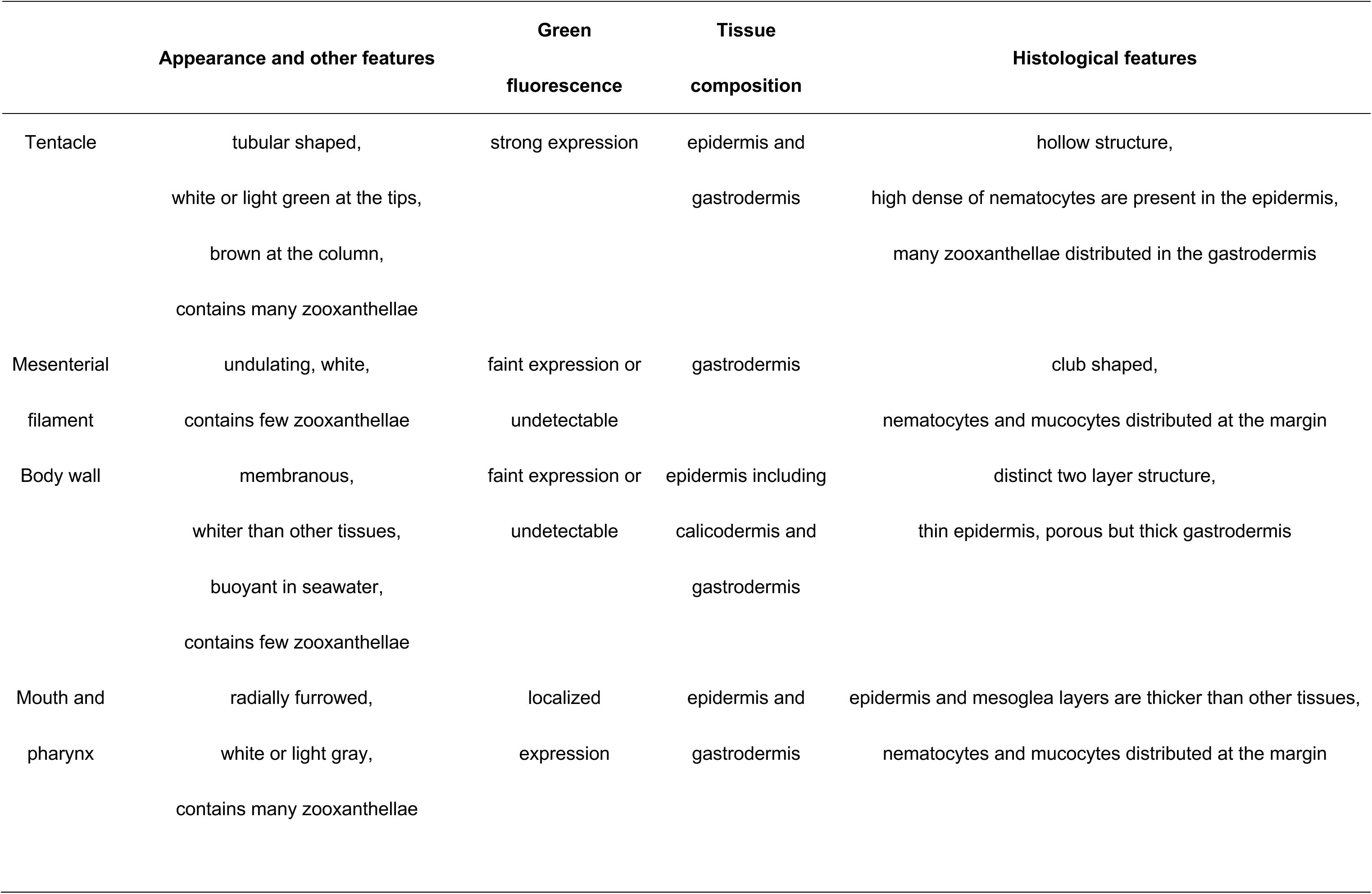
External and histological features of each tissue comprising *F. ancora polyps*.

|  | **Appearance and other features** | **Green**<br>**fluorescence** | **Tissue**<br>**composition** | **Histological features** |
| --- | --- | --- | --- | --- |
| Tentacle | tubular shaped,<br>white or light green at the tips,<br>brown at the column,<br>contains many zooxanthellae | strong expression | epidermis and<br>gastrodermis | hollow structure,<br>high dense of nematocytes are present in the epidermis,<br>many zooxanthellae distributed in the gastrodermis |
| Mesenterial<br>filament | undulating, white,<br>contains few zooxanthellae | faint expression or<br>undetectable | gastrodermis | club shaped,<br>nematocytes and mucocytes distributed at the margin |
| Body wall | membranous,<br>whiter than other tissues,<br>buoyant in seawater,<br>contains few zooxanthellae | faint expression or<br>undetectable | epidermis including<br>calicodermis and<br>gastrodermis | distinct two layer structure,<br>thin epidermis, porous but thick gastrodermis |
| Mouth and<br>pharynx | radially furrowed,<br>white or light gray,<br>contains many zooxanthellae | localized<br>expression | epidermis and<br>gastrodermis | epidermis and mesoglea layers are thicker than other tissues,<br>nematocytes and mucocytes distributed at the margin |

#### Tentacle

Tentacles were basically tubular, and the tips were hammer- or anchor-like **(Fig. 2 A-C)**. They were hollow and expanded by retaining seawater inside. Each tentacle was 3-5 cm long and 1-2 cm wide when fully expanded. Tips were white to light green, and the columns were brown, due to zooxanthellae **(Fig. 2 B)**. The white area at the tip showed strong green fluorescence under a fluorescence microscope **(Fig. 2 D and E)**. Histologically, a distinct two-cell layer, epidermis and gastrodermis, was observed. A high density of nematocytes and mucocytes was present in the epidermis, whereas gastrodermis cells and zooxanthellae were observed in the gastrodermis **(Fig. 2 F)**.

#### Mesenterial filament

Mesenterial filaments, located in the aboral part of the polyp, are formed by gastrodermis extending toward the body cavity **(Fig. 2 A and C)**. Isolated mesenterial filaments were white to pale yellow with an undulating structure **(Fig. 2 G)**. Individual mesenterial filaments (when contracted) were 2-3 mm in diameter. Slight green fluorescence was observed **(Fig. 2 H)**. Histologically, the margins were distinctive and club-shaped, and only a few zooxanthellae were observed. A number of nematocytes and mucocytes were present in the marginal area **(Fig. 2 I)**.

#### Body wall

The isolated part of the body wall was white **(Fig. 2 J)** and buoyant in seawater. Little green fluorescence was observed under the fluorescence microscope **(Fig. 2 K)**. Histologically, the body wall consisted of a distinct two-cell layer, a thin epidermis containing a calicodermis layer, and a relatively thick, porous gastrodermis **(Fig. 2 L)**. Only a few nematocytes, mucocytes, and zooxanthellae were observed in the body wall.

#### Mouth with pharynx

The isolated piece of mouth and pharynx was white to gray with a furrowed structure **(Fig. 2 M)**. Under the fluorescence microscope, spatial localization of green fluorescence was observed in the mouth, but not in the pharynx **(Fig. 2 N)**. Histologically, similar to tentacles, the mouth and pharynx consisted of two distinct layers, the epidermis with many nematocytes and mucocytes and the gastrodermis with many zooxanthellae **(Fig. 2 O).**

### Transcriptome analysis of the four tissues

Numbers of genes expressed in tentacles, mesenterial filaments, the body wall, and the mouth with pharynx were 19057, 19582, 18777, and 19587, respectively **(Fig. 3 A)**. Hierarchical clustering and a correlation heatmap showed that tentacles have the most specialized expression profiles **(Fig. 3 B)**. NMDS analysis further showed that transcriptional profiles of the four tissues were significantly different **(Fig. 3 C)**. In order to identify genes that characterize each tissue, we examined tissue-specific, highly expressed genes (HEGs): tentacles (856) mesenterial filaments (326), body wall (479), and mouth and pharynx (173) (q-value< 0.05, **Fig. 3 D**). Functional enrichment analysis of HEGs (**Fig. 3 E**) revealed tentacle-specific HEGs related to neurotransmitter transport, amino-acid transport, chloride channels, toxins, and nematocysts (> 6-fold enrichment). Mesenterial filament-specific HEGs were related to polysaccharide degradation, serine proteases, and protease inhibitors, whereas body wall-specific HEGs were associated with basement membrane and extracellular matrix. In mouth and pharynx, genes associated with the Wnt signaling pathway were significantly enriched **(Fig. 3 E).**

**Figure 3.**
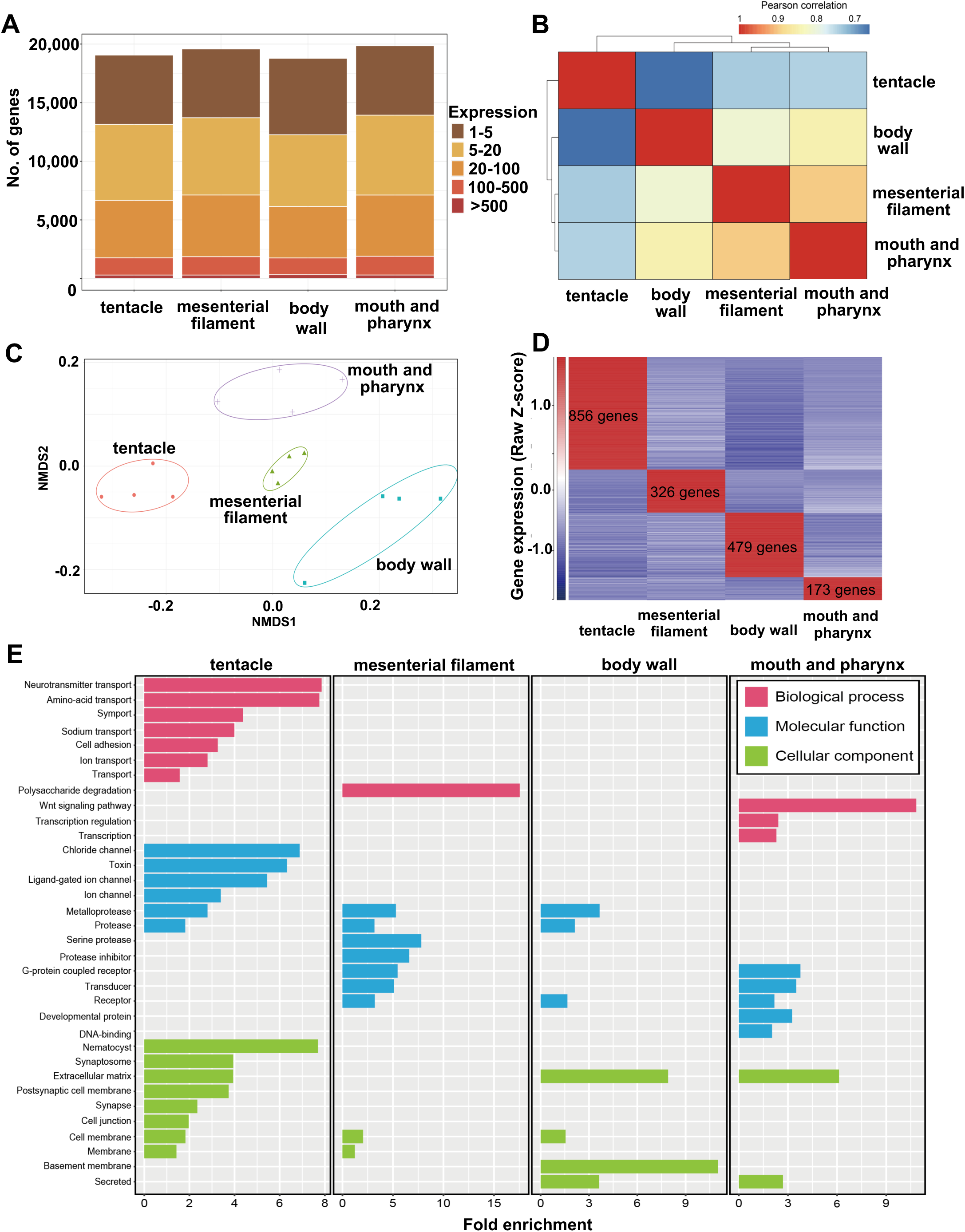
Classification of all protein-coding genes with regard to transcriptional levels in the 4 polyp tissues. **A**. Total numbers of genes with detected transcripts in each tissue type using five different abundance levels for transcripts per million (TPM) values; 1–5 TPM (brown), 5–20 TPM (dark yellow), 20–100 TPM (light orange), 100–500 TPM (dark orange), more than 500 TPM (dark red). 19057, 19582, 18777, and 19587 genes were detected in tentacle, mesenterial filament, body wall, and mouth and pharynx, respectively. **B**. Pearson correlation of tissue-specific transcriptomes of the 4 tissues using 16,703 expressed genes in all samples (TPM, >1). **C.** Nonmetric multidimensional scaling (NMDS) plot based on Bray-Curtis distances of gene expression profiles from tentacle, mesenterial filament, body wall, and mouth and pharynx. 12,302 genes TPM > 1 in 16 samples) were used for comparisons. Colored lines surrounding each sample type represent covariance ellipsoids. **D**. Relative gene expression levels of specifically highly expressed genes (in total 1834 genes) in the 4 polyp tissues. TPM values were scaled to row Z-scores for each gene. **E**. Significantly upregulated UniProt keywords detected from tissue-specific HEGs. Significantly (> 4- fold change, P < 0.05) enriched UniProt keywords in biological processes (red bar), molecular function (blue bar), and cellular component (green bar). The X-axis represents the magnitude of fold enrichment. The Y-axis represents the functional category.

### Biological characters of the four tissues revealed by transcriptomic analysis

Based on results of morpho-histological observation and functional enrichment analysis of HEGs described above, we explored and further selected expressed genes characterizing each tissue.

#### Tentacle

Tentacles highly expressed genes associated with biological defense and predation **(Fig. 4)**. This included cnidocytes (*Nematocyst expressed proteins*), venom (*DELTA-thalatoxin, DELTA-alicitoxin, and U-actitoxin*), mucus (*Mucin-like protein*), innate immunity (*Techylectin-5B*, *Antimicrobial peptide*, *Toll-like receptor 2*, and *Scavenger receptor*), and defense against oxidative stress (*Cell surface Cu-only superoxide*, *Glutathione peroxidase 5*, *Catalase*, *GFP-like fluorescent chromoprotein*). Tentacles also highly expressed genes encoding extracellular matrix (*Collagen alpha-1,2,4, and 5*, *Fibronectin*) and a cell adhesion molecule (*Protocadherin Fat 4*), which may support the three-dimensional structure of tentacles. Genes associated with coral-algal symbiosis/metabolic interactions (*Amino acid transporter AVT1A*, *Major facilitator superfamily domain-containing protein 10*, *NPC intracellular cholesterol transporter 2 homolog*, and *Ammonium transporter Rh type B-B*) were also detected. Moreover, genes for the light-sensing molecule, melanopsin-B, and the precursor of Antho-RF amide neuropeptide, which is involved in tentacle contraction, were also highly expressed. Two genes encoding a steroidogenesis-related enzyme, Steroid 17-alpha-hydroxylase/17,20 lyase, were also highly expressed **(Fig. 4).**

**Figure 4.**
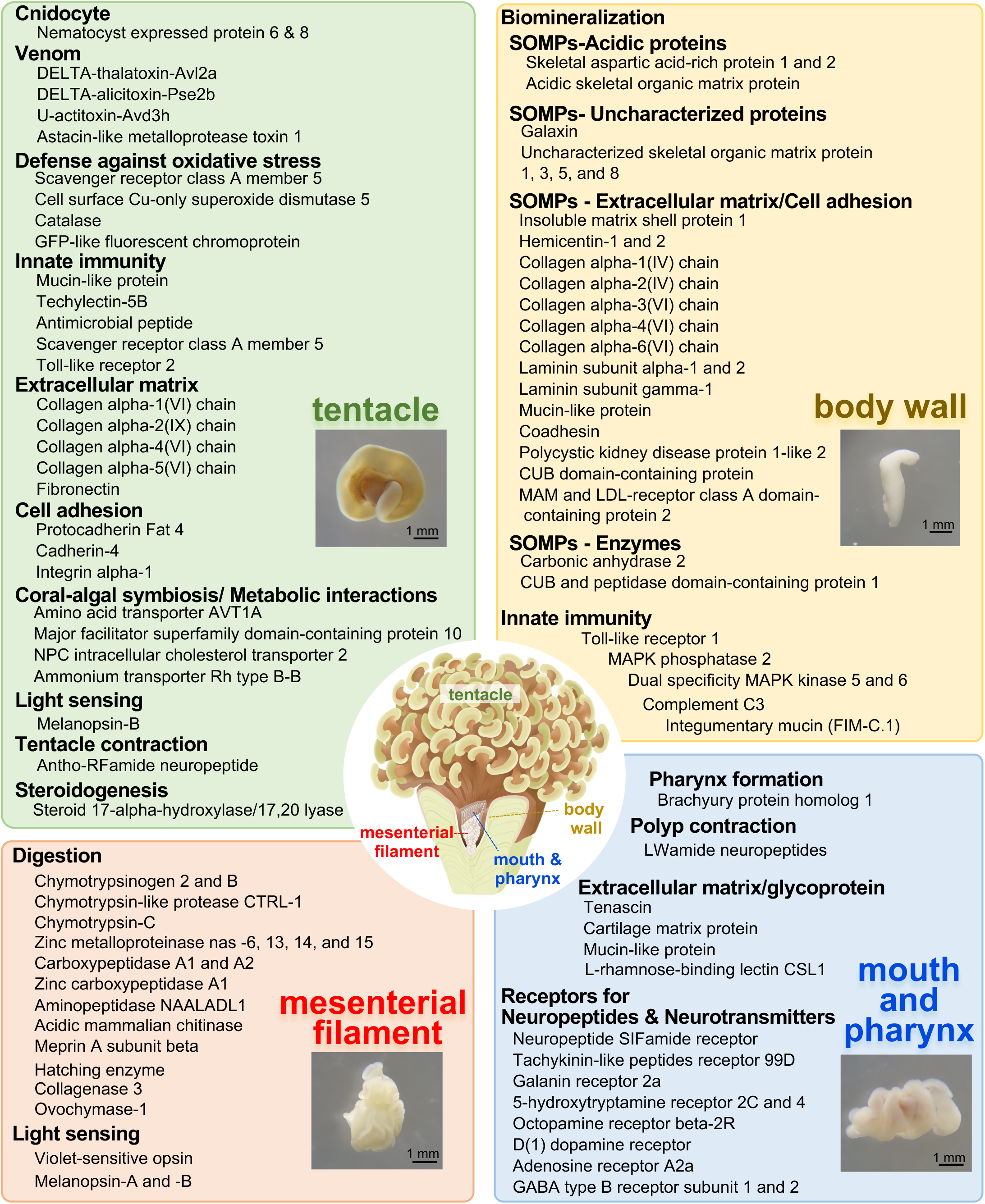
Possible functions of the 4 major tissues comprising *F. ancora* polyps and their potentially associated proteins. The micrograph in each panel shows isolated tentacle (left top), mesenterial filament (left bottom), body wall (right top), and mouth and pharynx (right bottom).

#### Mesenterial filament

Mesenterial filaments highly expressed a number of genes encoding proteases such as chymotrypsinogens, chymotrypsins, zinc metalloproteinases, carbopeptidases, aminopeptidase, meprin A subunit beta, and acidic mammalian chitinase. Mesenterial filaments also highly expressed genes encoding light-sensing molecules such as violet-sensitive opsin, melanopsin A, and melanopsin B **(Fig. 4)**.

#### Body wall

Body wall highly expressed a number of genes encoding skeletal organic matrix proteins (SOMPs), e.g., acidic SOMPs, uncharacterized protein, extracellular matrix, cell adhesion molecules, and enzymes **(Fig. 4)**. Genes encoding molecules related to innate immunity (Toll-like receptor 1, MAPK 2/5/6, Complement C3, integumentary mucin) were also highly expressed **(Fig. 4)**.

#### Mouth and pharynx

In mouth and pharynx, genes encoding a transcription factor, Brachyury, essential for pharynx formation during embryogenesis in corals, were highly expressed. Molecules related to the Wnt signaling pathway, innate immunity, and the precursor of GLWamide neuropeptides, involved in polyp contraction, were also detected **(Fig. 4).** Mouth and pharynx also highly expressed a number of receptors for neuropeptides, e.g., neuropeptide SIFamide, tachykinin-like peptides, and neurotransmitters, e.g., 5-hydroxytryptamine, octopamine, dopamine, GABA **(Fig. 4)**.

### Tissue-specific expression patterns of fluorescent protein, nk homeobox, and wnt genes

Coral color is primarily attributed to pigment proteins, including fluorescent proteins (FPs), non-fluorescent chromoproteins, and brown-pigmented zooxanthellae. In the *F. ancora* genome, we identified 11 candidate genes encoding FPs. Genes encoding non-fluorescent chromoproteins were not identified. Based on molecular phylogenetic analysis with two known *F. ancora* FP genes (one GreenFP and one RedFP) (Chiu et al., 2019; Shikina et al., 2016a) and FP-like genes from diverse anthozoan taxa, we estimated that 3 of the identified FP candidate genes encode GreenFPs (GFPs), while 1 encodes a RedFP (RFP) **(Fig. 5 A)**. Our molecular phylogenetic analysis also showed that 7 of 11 FP candidate genes in *F. ancora* were expanded by tandem duplication independently of those in *Acropora* **(Fig. 5A)**. These 7 FP genes are also arranged in tandem in the genome (**Fig. 5 B**). Tissue-specific transcriptome analyses revealed that 6 of 11 FP candidate genes, including 1 GFP gene, were highly and significantly expressed in tentacles **(Fig. 5 C)**, in agreement with fluorescent microscopy. One FP-like gene, s020 g104, is also highly, but not significantly, expressed in the mouth and pharynx **(Fig. 5 C)**. Immunohistochemical analysis with anti-GFP antibody showed that GFP was expressed in epidermis of tentacles **(Fig. 5 D)**.

**Figure 5.**
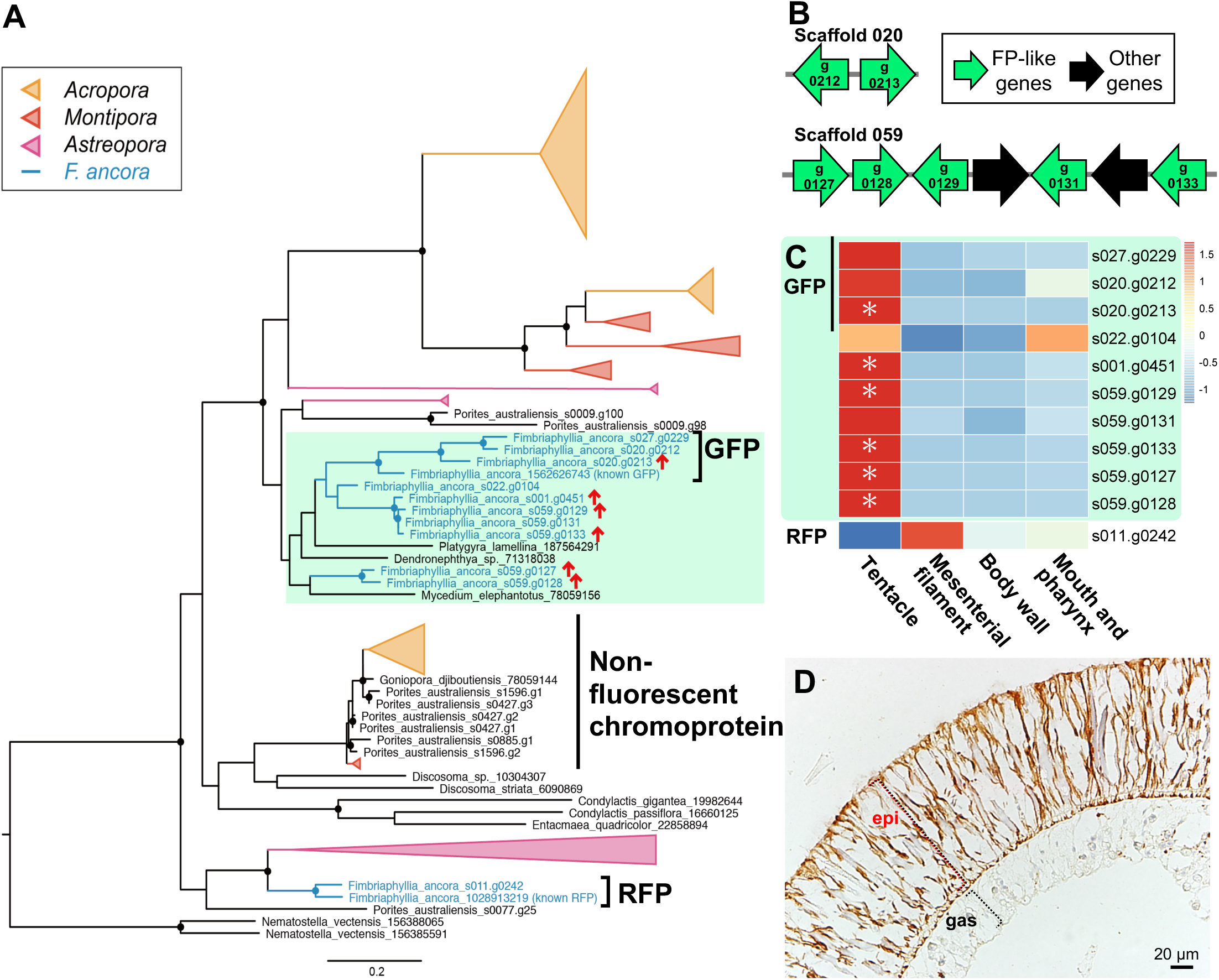
**A.** Molecular phylogenetic analysis of possible *F. ancora* FP-like genes with two known *F. ancora* FP genes (one GreenFP and one RedFP), FP genes, and non-fluorescent chromoprotein genes of diverse anthozoan taxa, including *Acropora*, *Montipora*, and *Astreopora* corals. Red arrows indicate tentacle-specific HEGs. Circles on branches indicate bootstrap values higher than 80%. **B**. Arrangement and orientation of 7 FP-like genes (shown as green arrows) in the *F. ancora* genome. Numbers in arrows indicate gene IDs in scaffolds. **C**. A heatmap showing transcript levels of 11 FP-like genes in tentacle, mesenterial filament, body wall, and mouth and pharynx. Asterisks indicate tissue-specific HEGs. **D**. Epidermal expression of GFP in the tentacle as assessed by immunohistochemical analysis with anti-GFP antibody.

Homeobox genes function in developmental patterning in animals (Hubert and Wellik, 2023). In the *F. ancora* genome, the Antennapedia (ANTP) class of homeobox genes, including Hox-like, ParaHox-like, and NK cluster genes, is present in the *F. ancora* genome. Hox and Hox-related gene clusters were found in scaffold 50, whereas two ParaHox genes were in scaffold 38 **(Fig. S1)**. Tissue distribution analysis found that one of the duplicated Mnx genes (g174) was more highly expressed in mesenterial filaments than in the other three tissues **(Fig. S1)**. NK cluster genes were found in scaffolds 12 and 19 **(Fig. 6 A)**. Tissue distribution analysis revealed that 8 of 10 NK genes cluster in scaffold 19 and all 4 in scaffold 12 showed tissue-specific expression patterns **(Fig. 6 A)**: 3 *MSX* genes in mouth and pharynx, two *NKX2* genes and one *NKX3* gene in body wall, *HLX, LBX, NKX3* and two *NKX2* genes in mesenterial filaments, and one *MSX* gene in tentacle **(Fig. 6 A and B)**.

**Figure 6.**
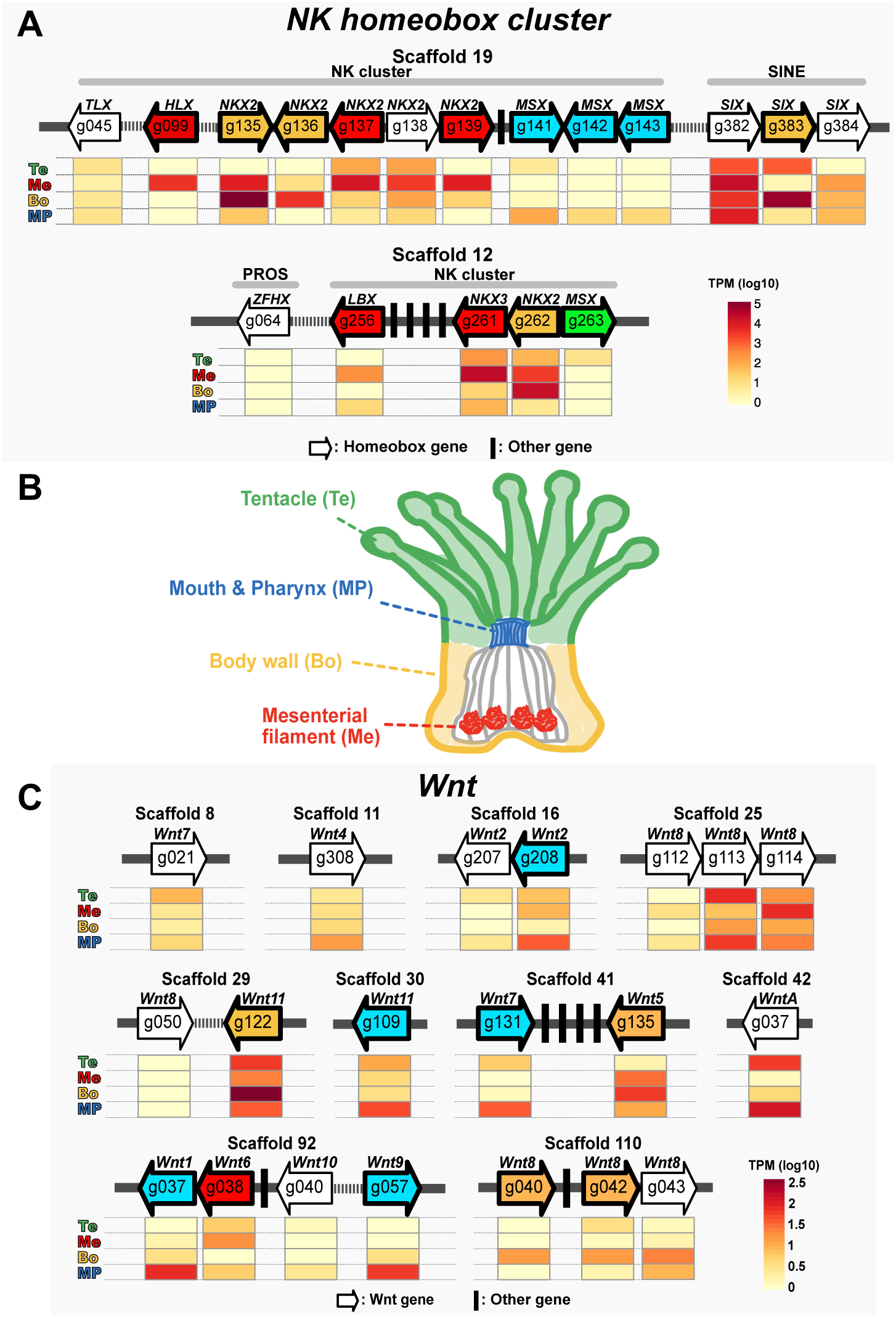
**A.** Organization of NK homeobox gene clusters in the *F. ancora* genome. Gene IDs and their relative positions and orientations in scaffolds 19 and 12 are indicated with arrows. Arrow directions indicate directions of transcription. Arrow colors indicate the tissue in which genes are highly expressed. Relative expression levels in the four polyp tissues are shown with heat maps under the arrows. Te, tentacle (green); MP, mouth and pharynx (blue); Bo, body wall (yellow); Me, mesenterial filament (red). **B**. Schematic diagram of a coral polyp showing locations of the 4 tissues analyzed in this study. **C**. Organization of Wnt genes in the *F. ancora* genome. Gene IDs and their relative positions and orientations in scaffolds are indicated by arrows. As in **A** above, colors of arrows indicate tissues in which genes are highly expressed. Relative expression levels in 4 tissues are shown with heat maps under the arrows.

The Wnt gene family encodes secreted signaling molecules that control cell fate in animal development and disease. In the *F. ancora* genome, 20 possible Wnt genes with conserved wnt family domains (PF00110) were identified (**Fig. 6 C, Fig. S2**) and the Wnt signaling pathway is enriched in mouth and pharynx, as mentioned above (Fig. 4). Among the 12 Wnt subfamilies (Wnt A, 1-11), an orthologous gene for Wnt3 was not identified (**Fig. 6 C**). Possible Wnt 8 genes were tandemly duplicated (3 genes in both scaffolds 25 and 110, respectively, **Fig. 6 C**). Tissue distribution analysis found that 10 of 20 Wnt genes were assigned as tissue-specific HEGs (**Fig. 6 C**): 5 genes (*Wnt1,-2,-7,-9*, and *-11*) in mouth and pharynx, 4 genes (*Wnt5,-8,-8* and *-11*) in body wall, and 1 gene (*Wnt6*) in mesenterial filaments **(Fig. 6 C)**

## Discussion

### Establishment of an F. ancora draft genome and the first tissue-specific gene expression profiles of coral polyps

First, we successfully established a draft genome of *F. ancora*. Comparison of this genome with those of other corals showed that the *F. ancora* genome has relatively high quality in terms of scaffold N50 length, numbers of scaffolds, and BUSCO completeness for genome assembly and gene models. Second, we established tissue-specific transcriptome assemblies of the four main tissues comprising *F. ancora* polyps. Previously, we established an *F. ancora* gonadal transcriptome assembly that allowed us to identify sex-and gonadal, phase-specific genes associated with germ cell development (Chiu et al., 2020). Combining this, tissue-specific transcriptomics of 6 tissues, i.e., tentacles, mesenterial filaments, body wall, mouth with pharynx, testes, and ovary, are now available for *F. ancora.* To the best of our knowledge, although genomes of more than 25 coral species have been reported so far (Shinzato et al., 2011; Voolstra et al., 2017; Cunning et al., 2018; Ying et al., 2019; Shinzato et al., 2021a; Shinzato et al., 2021b; Yoshioka et al., 2022), global classification of genes with regard to spatial expression patterns across tissues has not been undertaken in those species. Tissue-specific transcriptomic datasets of *F. ancora* in this study not only shed light on unexplored functions of coral tissues, but provide a useful foundation for better understanding of gene functions, as well as biological and physiological characteristics of corals.

### Tentacle: a multifunctional tissue involving biological defense, predation, external-factor sensing, symbiosis, and steroid synthesis

Our microscopic and histological observations demonstrated that some nematocytes and mucocytes are located in the epidermis, whereas zooxanthellae are present in the gastrodermis of tentacles. Consistent with these observations, genes associated with defense and predation, i.e., cnidocytes, venom, (Moran et al., 2013; Shiomi et al., 1997; Klompen et al., 2020; Trevisan-Silva et al., 2010) and defense against oxidative stress (Peskin, 1997; Gebicka and Krych-Madej, 2019) are highly expressed in tentacles. Genes involved in innate immunity (Sheng and Hasnain, 2022; Angthong et al., 2017; Mason et al., 2021; Areschoug and Gordon, 2009; Duan et al., 2022; Yoshioka et al., 2021) and light-sensing (Mason et al., 2012; Emanuel and Do, 2023) are also highly expressed. Furthermore, genes related to the three cellular mechanisms likely involved in symbiosis (antioxidant defense, immune regulation, and metabolite exchange) (Yoshioka et al., 2023) are also highly expressed. Some genes orthologous to symbiosis-related genes reported in the stony coral, *Acropora tenuis*, are also present (Yoshioka et al., 2023). These findings indicate that tentacle is a multifunctional tissue that not only serves as the front line of biological defense and predation, but also functions as a light-sensing tissue.

Many stony corals have the ability to produce diverse colored pigments. FPs, together with non-fluorescent pigment proteins (chromoproteins) and brown-pigmented zooxanthellae, create vivid displays of coral color (Dove et al., 2001; Alieva et al., 2008). While functions of FPs have yet to be discovered, since FPs absorb ultraviolet A and emit light with lower energy, proposed functions include photoprotection from high UVA/blue irradiation and photosynthetic enhancement of zooxanthellae (Salih et al., 2000; Roth et al., 2010). In addition, some studies suggest that FPs not only serve antioxidant functions (Bou-Abdallah et al., 2006; Palmer et al., 2009a; Shikina et al., 2016a), but also contribute to innate immunity (Palmer et al., 2009b; D’Angelo et al., 2012), stress response (Rodriguez-Lanetty et al., 2009; Seneca et al., 2010), and prey attraction (Ben-Zvi et al., 2022). In our previous study, we reported identification of RedFP (RFP) gene, specifically expressed in oocytes of *F. ancora*, and suggested its involvement in coral oogenesis (Shikina et al., 2016a). In the present study, *F. ancora* genomic and tissue-specific transcriptome analyses identified 6 FP genes that are highly upregulated in tentacles. This finding was also supported by fluorescence microscopic observations. Our phylogenetic analysis suggested that multiple FP genes highly expressed in *F. ancora* tentacles must have originated by independent gene duplication. Although their functions are currently unclear, they may have acquired functions specific to tentacles during evolution of the *F. ancora* lineage. FPs expressed in the epidermis of *F. ancora* tentacles may be involved in protecting coral cells and zooxanthellae in the polyp from intense UV light, or in attracting prey.

Of particular interest is the significantly higher expression of the gene encoding steroid 17α-hydroxylase/17,20-lyase (Cyp17a), a key enzyme in production of sex steroids and cortisol (Guengerich et al., 2023), in *F. ancora* tentacles. Steroids are biologically active compounds derived from cyclopenta[a]phenanthrene (Sultan and Rauf-Raza, 2015). They function as important components of cell membranes and are involved in a wide range of physiological processes, such as stress response, immune response, behavior, reproduction, etc. (Rasheed and Qasim, 2013). Steroid biosynthesis is catalyzed by activities of various steroidogenic enzymes (Miller and Auchus, 2011). To date, steroidogenic enzyme activities, plus estrogens and testosterone have been demonstrated in tissue extracts of some scleractinians (Atkinson and Atkinson, 1999; Tarrant et al., 1999; Slattery et al., 1999; Twan et al., 2003). Further, some genes encoding steroidogenic enzymes were also demonstrated (Shikina et al., 2016b; Tan et al., 2021). Although further research is required to clarify transcript localization and steroidal activity, our findings imply that steroid biosynthesis occurs in tentacles, and that produced steroids/cortisol could be associated with important functions in tentacles, such as maintenance of cell membrane integrity, biological defense, or symbiosis with zooxanthellae.

### Digestion and light-sensing in mesenterial filaments

Our histological analysis demonstrated that many cnidocytes and mucocytes are present in the margins of mesenterial filaments. Further, we identified a variety of digestive enzymes that are highly expressed in mesenterial filaments. Since corals feed on various types of prey (Poter, 1974; Seben et al., 1996), it is possible that expression levels of the identified digestive enzymes may also vary depending on types of prey. Nevertheless, coral mesenterial filaments have so far been studied mainly from a behavioral perspective (Nugues et al., 2004, Roff et al., 2009). There is only one paper describing gene expression (Raz-Bahat et al., 2017).

Some coral behaviors such as tentacle expansion/contraction are strongly influenced by light (Abe, 1939; Sweeney, 1976; Sebens and Deriemer, 1977; Lasker, 1979; Hoadley et al., 2011). *F. ancora* also extends its polyps in response to light (data not shown). Additionally, it usually releases gametes at night (Shikina et al., 2023b). However, it is still largely unknown how corals, which have no special light-sensing organs, perceive light. Opsin and melanopsins, key molecules in photosensing (Emanuel and Do, 2023), exhibit tissue-specific expression in *F. ancora*. There are 32 genes encoding opsins/melanopsins in the *F. ancora* genome. Melanopsin B is highly expressed in tentacles, whereas Violet-sensitive opsin, Melanopsin A, and Melanopsin B are highly expressed in mesenterial filaments. We hypothesize that corals may enhance sensitivity to light by highly expressing photo-sensing molecules not only in tentacles, but also in mesenterial filaments. In *Acropora millepora*, multiple opsins have been identified (Mason et al., 2012), including an opsin that responds to UV light (Mason et al., 2023).

### Body wall: skeleton formation and chemical-physical barrier

Nematocytes and zooxanthellae are scarcely present in the body wall of *F. ancora*, suggesting that body wall is basically not involved in predation or symbiosis. Surprisingly, body wall highly expresses a variety of molecules involved in biomineralization (acidic SOMPs, uncharacterized proteins, extracellular matrix, cell adhesion molecules, and enzymes) (Ramos-Silva et al., 2013; Drake et al., 2013; Takeuchi et al., 2016). This is possibly because *F. ancora* forms its skeleton entirely outside the polyp body wall. The epidermis of the body wall basically serves as a layer of calicodermis, in contact with the skeleton. Further, the body wall also expresses some molecules involved in innate immunity, such as Toll-like receptor 1, Complement C3, integumentary mucin (Dunkelberger and Song, 2010; Duan et al., 2022; Sheng and Hasnain, 2022). These results indicate that the body wall is not only the primary tissue for the skeleton formation, but also functions as a chemical-physical barrier to parasitic and biofouling organisms.

### Mouth and pharynx: key tissues for coral nervous system?

A notable finding was the high expression of many neuropeptides and biogenic amine receptors in the mouth and pharynx. Both peptidergic and non-peptidergic neurotransmission/neuromodulation are fundamental to cnidarian physiology (Kass-Simon and Pierobon, 2007; Takahashi and Takeda, 2015; Takahashi, 2020). To date, in corals, GLWamide-positive and RFamide-positive neurons have been found in the mouth and pharynx. Both of these are involved in polyp contraction (Attenborough et al., 2019; Shikina et al., 2020, Zhang et al., 2021). Some neuroactive compounds such as dopamine, glutamic acid, and epinephrine induce larval settlement in the coral, *Leptastrea purpurea* (Moeller et al., 2019). Increases in levels of dopamine, adrenaline, and noradrenaline were detected during a synchronous spawning event in *Acropora intermedia* (Taira et al., 2018). Evidence for neuropeptides and neurotransmitters in coral physiology is slowly accumulating; however, much of it remains unknown. Many coral neuropeptides, neurotransmitters, and their receptors have not yet been identified. Their identification will provide a more complete picture of coral nervous systems. Future investigations will employ liquid chromatography-tandem mass spectrometry (LC-MS/MS) and biotin-tagged neuropeptides with LC-matrix-assisted laser desorption/ionization (MALDI)-MS/MS (Shinya et al., 2010), or recent machine learning-associated research strategies (Shiraishi et al., 2019; Satake et al., 2023), targeting especially the mouth and pharynx.

### Possible involvement of NK homeobox clusters and Wnt genes in establishment of tissue-specific gene expression in coral polyps

How are the distinct gene expression patterns and unique biological characteristics of the four tissues governed and maintained? The presence of homeobox genes, including Hox-like and ParaHox-like, and NK cluster genes, was demonstrated in the *F. ancora* genome, as reported in some other corals (DuBuc et al., 2012; Ying et al., 2018), but only one Mnx gene (eanc_s050.g174) in the Hox-like gene cluster (10 genes) was assigned as a tissue-specific HEG (Supp. Fig. S1). Notably, tissue distribution analysis found that 11 of 14 genes in NK homeobox clusters showed tissue-specific expression patterns in *F. ancora* polyps. In a sea anemone, *Nematostella vectensis* (Anthozoa), NK homeobox genes show tissue-specific expression patterns. *Gbx* is expressed in pharyngeal endoderm (Matus, et al. 2006), *Hlx* and *Nk6* in pharyngeal ectoderm, and *Nk3* in nutrient-storing somatic gonads in mesentery (Steinmetz, et al. 2017). Although anthozoans possess a pharynx and a mesentery, medusozoans do not have these, possibly reflecting extensive loss of NK genes and fragmentation of the NK cluster (Hamada, et al. 2020). NK cluster genes may function in body plan patterning and underlying tissue specificity in anthozoans. Of particular interest is the discovery of significantly higher expression of genes encoding various types of Wnt protein in *F. ancora* mouth and pharynx. Wnt signaling pathways are found exclusively in animals and contribute to development and cell differentiation even in cnidarians, the basal animal phylum (Loh et al., 2016). In *Hydra*, various types of Wnt mRNA (*HyWnt1, −3, −7, −9/10a*, *-9/10c, −11,* and *-16*) are expressed in hypostomes of both adult polyps and new buds, and their involvement in head formation was demonstrated (Hobmayer et al., 2000; Lengfeld et al., 2009). In *Nematostella*, induction of Wnt signaling with alsterpaullone results in formation of ectopic oral tissue during regeneration and embryogenesis (Trevino et al., 2011). Accordingly, Wnt may also be involved in formation of oral tissue in corals.

### Conclusions and Future Perspectives

Given the serious plight of coral reefs, promoting coral conservation, and increasing our understanding of coral biology is essential. Due to various unrelated tissues, investigations of whole coral polyps obscure important phenomena involved in coral beaching, breakdown of mutualistic symbiosis, and calcification, which supports the most biodiverse marine environments. As shown here, analyses focused on tissues involved in specific biological phenomena is essential to uncover physiological mechanisms. We hope that genomic information from *F. ancora* will facilitate coral conservation activities. Genome-wide SNP markers will allow us to reveal detailed population structures in nature (Shinzato et al., 2015; Tuchiya et al., 2022; Zhang et al., 2022), and will be applicable to *F. ancora* aquaculture to monitor genetic diversity in captivity (Shinzato et al., 2014; Zayasu et al., 2016; Zayasu et al., 2018; Zayasu and Suzuki, 2019). Although morpho-histological and transcriptome characteristics of coral polyp tissues are presented here, the number of cell types and compositional ratios of those cells constituting each tissue are still unknown. Single-cell transcriptome analysis (Levy et al., 2021) will clarify these issues.

## Materials and Methods

### Sampling, DNA and RNA extraction, transcriptome and genome sequencing

Four colonies of *F. ancora* were collected on reef slopes in the nearshore of Onna Village, Okinawa, Japan, under Okinawa prefectural permit #30-8 in 2018. Four polyp tissues, tentacle, mouth, mesenterial filament, and body wall, were isolated and photographed under a stereomicroscope (Shikina et al., 2013). Isolated tissues were snap frozen in liquid nitrogen and stored at −80 °C until use. Total RNA was isolated from each tissue with an RNeasy plant kit (QIAGEN Inc., Valencia, CA). For transcriptome sequencing, a TruSeq Stranded mRNA Library Kit (Illumina, San Diego, CA) was used for mRNA sequencing and library preparation, and each library was sequenced from 150-bp paired-end reads using a HiSeq 4000 (Illumina). To prepare genomic DNA for sequencing, isolated mesenterial filaments from one colony were maintained in Petri dishes for 1 week with 50 mL of filtered seawater containing 0.2 mM menthol (Wang et al., 2012) to eliminate symbiotic algae. Bleached mesenterial filaments were snap-frozen in liquid nitrogen and stored at −80 °C until use. Genomic DNA was isolated using the phenol-chloroform method, and sequenced on a PacBio platform to obtain highly accurate long-read sequencing data. Genome shotgun sequencing (150-bp paired-end) of the same DNA sample was also performed on an Illumina HiSeq4000 platform.

### Genome assembly and gene prediction

PacBio HiFi reads with quality values >20 were assembled with Hifiasm version 0.14-r312 using default settings (Cheng et al., 2021). Possible diploid scaffolds were first removed with Purge_haplotigs ver. 1.1.1 (Roach et al., 2018), and then merged with HaploMerger2 (Huang et al., 2017). Then further scaffolding was performed with LINKS (ver. 1.8.7) with a kmer size of 21 (Warren et al., 2015). Possible errors in genome assembly were corrected with Hypo ver. 1.0.3 (Kundu et al., 2019) using Illumina shotgun data with default settings. For gene prediction from assembled genome sequences, we used the above-prepared RNA-Seq data from the four tissues and RNA-Seq data from *F. ancora* male and female gonads in different developmental stages (Chiu et al., 2020). Low-quality reads (quality score < 20 and length < 20 bp) and sequence adaptors in RNA-Seq data were trimmed using CUTADAPT v1.18 (Martin, 2011). Repetitive elements in scaffolds were identified de novo with RepeatScout v1.0.6 (Price et al., 2005) and RepeatMasker v4.1.0 ( http://www.repeatmasker.org). Repetitive elements were filtered out by length (>50 bp) and occurrence. Gene prediction was first performed with the BRAKER pipeline v2.1.2 (Brůna et al., 2021), with AUGUSTUS v3.3.3. RNA-seq reads were aligned to each genome sequence with HISAT v2.1.0 (Kim et al., 2015). Then, alignment information was used for BRAKER gene prediction with options “UTR=on”, “soft-masking”, and “AUGUSTUS_ab_initio.” To improve gene prediction, we further executed genome-guided transcriptome assembly using StringTie (Pertea et al., 2015) with option “-m 500.” Genome-based transcript structure was predicted with TransDecoder (https://github.com/TransDecoder/ TransDecoder/wiki). During read alignment, we used soft-masked repeats for genome-guided transcriptome assembly and hard-masked repeats for BRAKER gene prediction. Finally, genes present in genome-guided assembly or ab initio prediction, but absent in predictions from the hint file were added to the predicted file using GffCompare (Pertea and Pertea, 2020). We assessed completeness of the genome assembly and gene prediction with Benchmarking Universal Single-Copy Orthologs (BUSCO) ver. 3.0.2 (Simão et al., 2015; Waterhouse et al., 2018) using the Metazoa set (978 genes).

### Gene annotation, clustering orthologous genes, and molecular phylogenetic analysis

Predicted gene models were BLASTed against the Uniprot/Swissprot (UniProt Consortium 2018) database and were analyzed with InterProScan 5 with cutoff of e^-5^ (Jones et al., 2014). For clustering orthologous genes in scleractinian genomes, we used four species of Acroporidae (*A. millepora*, *A. tenuis*, *Montipora cactus,* and *Astreopora myriophthalma*) (Shinzato et al., 2011; Ying et al., 2019; Shinzato et al., 2021a; Yoshioka et al., 2022), *Porites australiensis* (Shinzato et al., 2021b), *Stylophora pistillata* (Voolstra et al., 2017), *Pocillopora damicornis* (Cunning et al., 2018) and *Orbicella faveolata* (Prada et al., 2016). For the *A. millepora*, *S. pistillata*, *O. faveolata*, and *P. damicornis* genomes, we downloaded data from the NCBI RefSeq database. Then, using OrthoFinder version 2.5.4 (Emms and Kelly, 2015), we performed clustering of possible orthologs, and Orthogroups (OGs) were used for subsequent analyses.

For phylogenomic analysis of scleractinian genomes, we used 4,208 genes that were identified by OrthoFinder as single-copy genes in all of the above scleractinian genomes. All amino acid sequences belonging to same OG were aligned with MAFFT (ver. 7.310. with –auto option) (Katoh and Standley, 2013) and all gaps in the alignment were removed with TrimAL (Capella-Gutierrez et al., 2009) with the –nogaps option. Then all sequences from the same species were concatenated. Finally, a maximum likelihood analysis was performed using concatenated sequences (1,817,638 amino acids in length) from RAxML (maximum likelihood method) with 100 bootstraps replicates and the “protgammaauto” option (Stamatakis, 2014). For phylogenetic analysis of each gene, amino acid sequences were aligned using MAFFT (ver. 7.310. with –auto option) (Katoh and Standley, 2013), and gaps in aligned sequences were trimmed using TrimAL (Capella-Gutierrez et al., 2009) with the – gappyout option. After that, poorly aligned sequences were removed (-resoverlap 0.75 -seqoverlap 80). Then we performed molecular phylogenetic analysis of the selected alignments using RAxML (maximum likelihood method) with 100 bootstrap replicates and “protgammaauto” option.

### Tissue-specific gene expression analysis

RNA-Seq data obtained from the four tissues were used. Low-quality reads (quality score <20 and length <20 bp) and Illumina sequence adaptors were trimmed with CUTADAPT v1.16 (Martin 2011), and then mapped to *F. ancora* gene models (mRNA) using KALLISTO v0.44.0 (Bray et al., 2016) with 100 bootstrap replicates. Transcript abundances in each sample were quantified using SALMON v1.0.0 Mapping counts were normalized by the trimmed mean of M values (TMM) method, and then converted to counts per million (CPM) using EdgeR v3.32.1 in R v4.0.3. Gene expression levels (numbers of mapped reads) of each tissue were compared pairwise with the other 3 tissues. *p*-values were adjusted using the Benjamini– Hochberg method in EdgeR. When the gene expression level was significantly higher (False discovery rate < 0.05) in one tissue than the other three, genes were considered tissue-specific HEGs.

### Histological analysis

Histological analysis was performed according to the methodology described in our previous study (Shikina et al., 2012). Briefly, isolated tissues were fixed in filtered seawater containing 20% Zinc Formal Fixx (Thermo Scientific Shandon, Cheshire, UK) for 16 h and preserved in 70% ethanol until use. Dehydrated samples were embedded in paraplast plus (Sherwood Medical, St. Louis, MO), sliced into 4-mm serial sections, and stained with haematoxylin and eosin Y (H & E staining, Thermo Shandon). Stained sections were observed and photographed under a BX51 microscope (Olympus, Tokyo, Japan).

## Ethics

Experiments were conducted in accordance with principles and procedures approved by the Institutional Animal Care and Use Committee of National Taiwan Ocean University.

## Author contributions

SS and CS conceptualized and designed the project. YZ collected the specimens. SS and T-CL dissected tissues and collected samples. CS and NS supervised and organized whole genome sequencing and RNA-Seq. MiK, MaK and MF prepared sequencing libraries and produced sequencing data. CS assembled genomes and YY performed gene predictions. SS, YY, Yi-LC, TU, EC, Y-CC, Yu-LC, TT, and CS performed data analyses. SS and CS wrote the manuscript.

## Acknowledgments

We thank all members of the DNA Sequencing Section at OIST for their enormous help in the project. We also thank Dr. Steven Aird for his great help in preparing the manuscript.

## Figure Legends

**Supplementary Figure 1**

**A.** Organization of the Hox-like gene cluster in the *F. ancora* genome. Gene IDs and their relative positions and orientations in scaffolds 50 and 38 are indicated with arrows. Arrow directions indicate directions of transcription. Relative expression levels of genes in the four polyp tissues are shown with heat maps under the arrows. Te, tentacle; MP, mouth and pharynx; Bo, body wall; Me, mesenterial filament.

**Supplementary Figure 2**

Molecular phylogeny of *F. ancora* Wnt-related genes with Wnt genes from various taxa used in Kusserow et al. (2005). The phylogenetic tree was constructed using the maximum likelihood method with aligned amino acid sequences (147 amino acids). Numbers at nodes represent bootstrap probabilities >70%. Species abbreviations: Ag, *Anopheles gambiae*, Bf, *Branchiostoma floridae*; Bm, *Bombyx mori*; Ce, *Caenorhabditis elegans*; Dm, *Drosophila melanogaster*; Hs, *Homo sapiens*; Hv, *Hydra vulgaris*; Nv, *Nematostella vectensis*; Pd, *Plathynereis dumerlii*; Pv, *Patella vulg*

