## Supplementary data for "A genome and tissue-specific transcriptomes of the large-polyp coral, *Fimbriaphyllia* (*Euphyllia*) *ancora*: Recipe for a coral polyp"

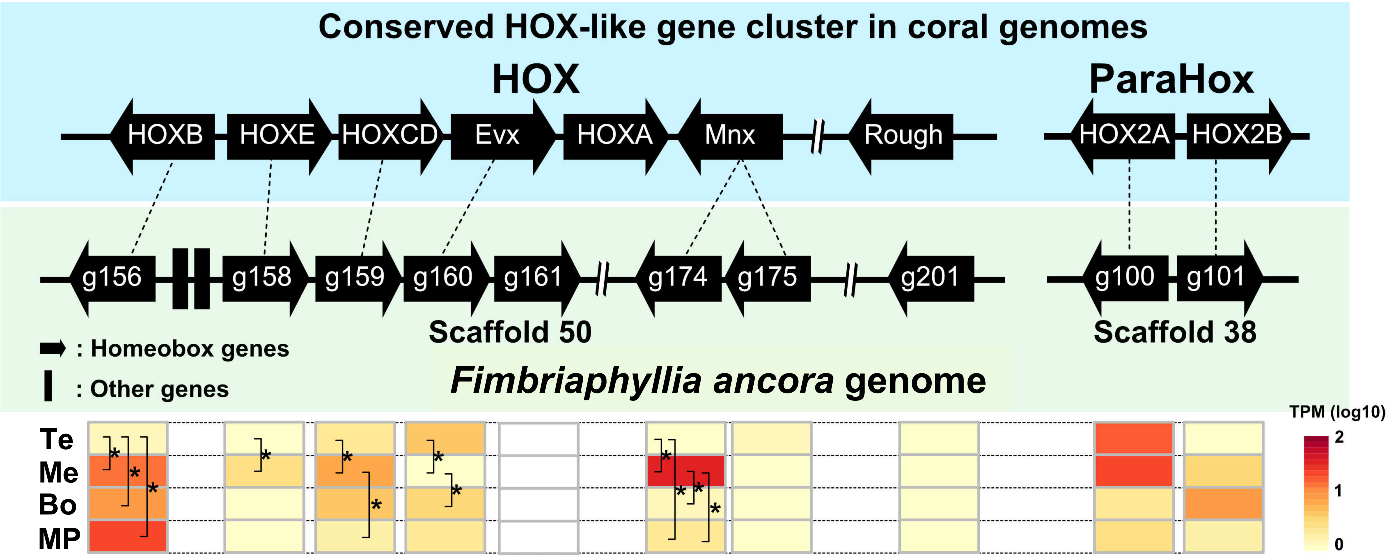
Conserved HOX-like gene cluster in coral genomes
HOX
ParaHox
HOXB
HOXE
HOXCD
Evx
HOXA
Mnx
Rough
HOX2A
HOX2B
g156
g158
g159
g160
g161
g174
g175
g201
g100
g101
Scaffold 50
Scaffold 38
: Homeobox genes
: Other genes
Fimbriaphyllia ancora genome
Te
Me
Bo
MP
TPM (log10)
2
1
0

**Fig. S1**

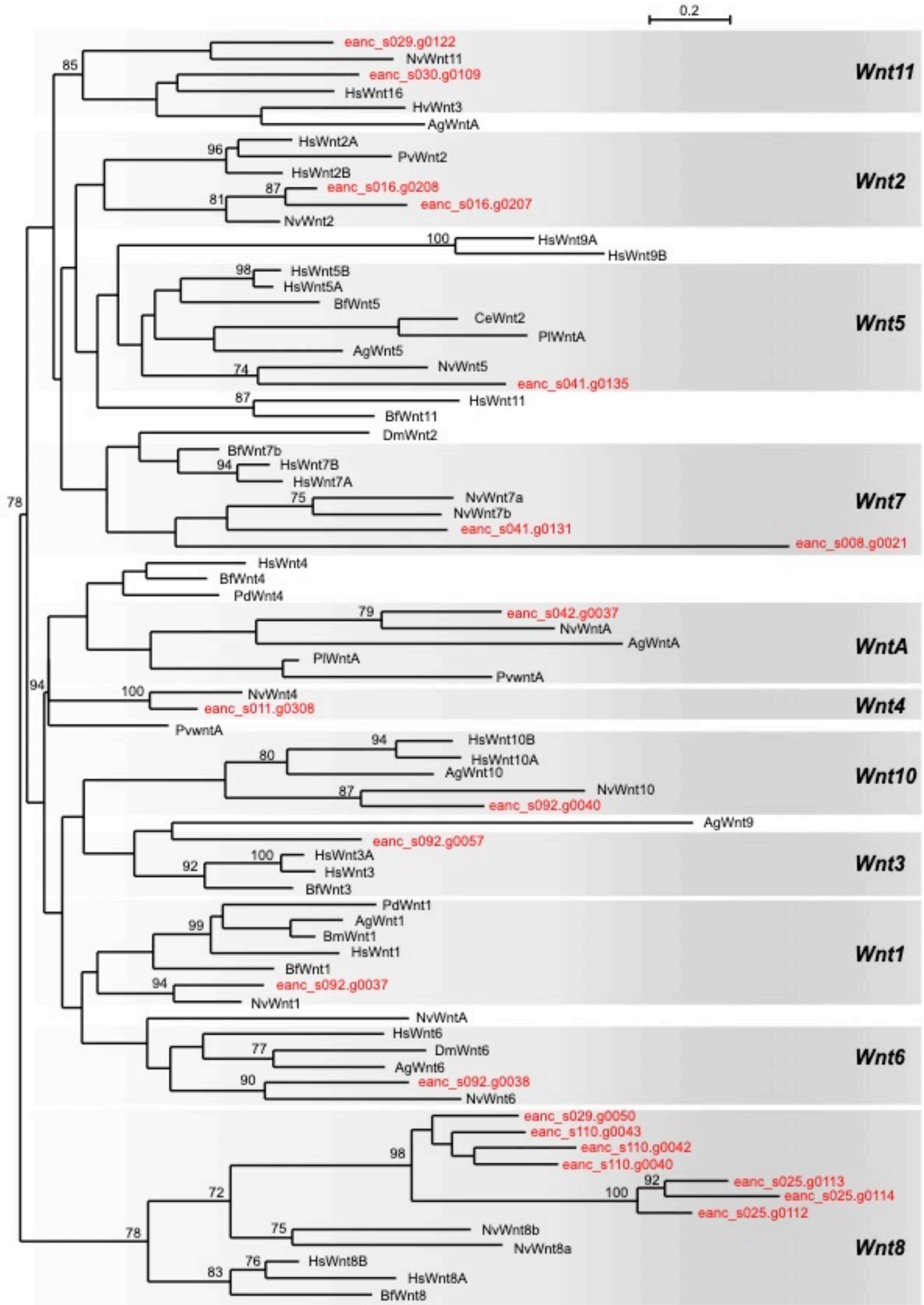
0.2
eanc_s029.g0122
NvWnt11
eanc_s030.g0109
HsWnt16
HvWnt3
AgWntA
Wnt11
HsWnt2A
PvWnt2
HsWnt2B
eanc_s016.g0208
eanc_s016.g0207
NvWnt2
Wnt2
HsWnt9A
HsWnt9B
HsWnt5B
HsWnt5A
BfWnt5
CeWnt2
PlWntA
AgWnt5
NvWnt5
eanc_s041.g0135
Wnt5
HsWnt11
BfWnt11
DmWnt2
BfWnt7b
HsWnt7B
HsWnt7A
NvWnt7a
NvWnt7b
eanc_s041.g0131
eanc_s008.g0021
Wnt7
HsWnt4
BfWnt4
PdWnt4
eanc_s042.g0037
NvWntA
AgWntA
PlWntA
PvwntA
WntA
NvWnt4
eanc_s011.g0308
Wnt4
PvwntA
HsWnt10B
HsWnt10A
AgWnt10
NvWnt10
eanc_s092.g0040
Wnt10
AgWnt9
eanc_s092.g0057
HsWnt3A
HsWnt3
BfWnt3
Wnt3
PdWnt1
AgWnt1
BmWnt1
HsWnt1
BfWnt1
eanc_s092.g0037
NvWnt1
Wnt1
NvWntA
HsWnt6
DmWnt6
AgWnt6
eanc_s092.g0038
NvWnt6
Wnt6
eanc_s029.g0050
eanc_s110.g0043
eanc_s110.g0042
eanc_s110.g0040
eanc_s025.g0113
eanc_s025.g0114
eanc_s025.g0112
NvWnt8b
NvWnt8a
HsWnt8B
HsWnt8A
BfWnt8
Wnt8

**Fig. S2**

**Table S1**. Results of homology search for the presence of Symbiodiniaceae-derived sequences within the established draft genome of *F. ancora*

| Scaffold # | GC content (%) | Hit length | Scaffold length | Ratio (%, Hit length / Scaffold length) |
|---|---|---|---|---|
| s001 | 39.66 | 257 | 16,539,788 | 0.00155383 |
| s002 | 39.65 | 886 | 12,320,501 | 0.00719127 |
| s003 | 39.76 | 778 | 11,729,092 | 0.00663308 |
| s004 | 39.6 | 75 | 10,172,913 | 0.00073725 |
| s005 | 39.63 | 301 | 10,071,468 | 0.00298864 |
| s008 | 39.7 | 553 | 8,583,838 | 0.00644234 |
| s009 | 39.59 | 183 | 10,227,658 | 0.00178927 |
| s010 | 39.65 | 224 | 8,226,613 | 0.00272287 |
| s012 | 39.74 | 85 | 7,167,972 | 0.00118583 |
| s013 | 40.02 | 839 | 12,098,484 | 0.00693475 |
| s014 | 39.45 | 218 | 6,970,475 | 0.00312748 |
| s015 | 39.6 | 221 | 6,716,729 | 0.00329029 |
| s017 | 39.34 | 217 | 6,106,193 | 0.00355377 |
| s019 | 38.97 | 84 | 5,904,139 | 0.00142273 |
| s021 | 40.06 | 259 | 6,199,459 | 0.00417778 |
| s024 | 39.82 | 603 | 7,413,012 | 0.00813435 |
| s029 | 39.05 | 83 | 4,809,300 | 0.00172582 |
| s031 | 39.25 | 93 | 4,183,812 | 0.00222285 |
| s033 | 39.52 | 76 | 4,155,996 | 0.00182868 |
| s037 | 39.66 | 77 | 3,571,058 | 0.00215622 |
| s048 | 39.81 | 206 | 3,441,960 | 0.00598496 |
| s049 | 39.65 | 778 | 5,519,598 | 0.01409523 |
| s055 | 39.81 | 131 | 2,586,332 | 0.00506509 |
| s059 | 39.54 | 249 | 2,483,856 | 0.01002474 |
| s063 | 39.69 | 123 | 2,635,939 | 0.00466627 |
| s064 | 39.57 | 305 | 2,235,725 | 0.01364211 |
| s070 | 39.33 | 674 | 1,850,587 | 0.03642088 |
| s071 | 39.8 | 78 | 1,839,249 | 0.00424086 |
| s090 | 39.93 | 362 | 1,418,153 | 0.02552616 |
| s091 | 39.28 | 135 | 1,226,400 | 0.01100783 |
| s106 | 39.39 | 99 | 897,672 | 0.01102853 |
| s112 | 42.93 | 48 | 796,304 | 0.00602785 |
| s115 | 39.05 | 319 | 771,582 | 0.04134363 |
| s132 | 40.75 | 778 | 440,772 | 0.17650849 |
| s140 | 41.07 | 86 | 335,880 | 0.02560438 |
| s152 | 39.18 | 82 | 346,997 | 0.02363133 |
